# Tumor emboli-associated adaptive stress response signatures identify aggressive disease features in inflammatory breast cancer

**DOI:** 10.64898/2026.07.06.734332

**Authors:** Pritha Pai, Hillary Hsu, Ganiraju C. Manyam, Steven Van Laere, David P. Mysona, William G. Hawkins, Savitri Krishnamurthy, Megumi Kai, Lixia Diao, Wendy A. Woodward, Gayathri R. Devi

## Abstract

Inflammatory breast cancer (IBC) is an aggressive breast cancer subtype characterized by tumor emboli, lymphovascular invasion (LVI), and early dissemination. Herein, we establish adaptive stress response (ASR) as a biologic feature linking stress adaptation to tumor emboli survival, lymphatic dissemination, therapeutic response, and disparities. Using a previously defined 226 ASR-related genes, complementary preclinical models of tumor emboli and lymphatic circulating cell clusters, and independent patient cohorts, we identified ASR genes enriched for XIAP-NFκB, oxidative stress response, inflammatory, and immune pathways. *CXCL8* emerged as one of the most highly upregulated transcripts in tumor emboli and was shared across both models; however, *CXCL8*, *IL6*, and *PTGS2* were downregulated in lymphatic circulating cell clusters and LVI-positive triple-negative IBC patients, suggesting dynamic remodeling of inflammatory signaling during dissemination. *CYP4B1* was associated with ER status, LVI, and therapeutic response across multiple cohorts, implicating metabolic stress adaptation in dissemination. *IL6* and *PTGS2* were elevated in self-reported Black patients with triple-negative IBC compared to White patients. Pharmacologic inhibition of XIAP-NFκB and oxidative stress pathways suppressed tumor emboli formation. Collectively, these findings identify ASR signaling as a framework linking tumor emboli survival, dissemination, and therapeutic vulnerability in IBC.

## Introduction

Inflammatory breast cancer (IBC) is a rare but highly aggressive form of breast cancer characterized by rapid progression, early metastatic dissemination, and poor clinical outcomes^1–3^ Unlike other breast cancer subtypes, IBC frequently presents without a dominant palpable mass and is instead defined by diffuse tumor cell clusters that invade breast and dermal lymphatic and blood vessels and form tumor emboli^4–7^. These multicellular emboli represent a histopathologic hallmark of IBC and are thought to play a central role in lymphatic route of invasion, early metastasis and therapeutic resistance^5,8,9^. Despite their clinical importance, the molecular mechanisms that enable tumor emboli to survive and persist within the hostile lymphovascular microenvironment remain poorly understood.

During dissemination, tumor emboli are exposed to multiple forms of cellular stress, including oxidative stress, inflammatory signaling, immune surveillance, and fluctuations in oxygen and nutrient availability^4,10–14^. To survive under these conditions, tumor cells activate adaptive stress response (ASR) pathways that suppress programmed cell death and promote cellular survival and phenotypic plasticity^15–17^. In cancer, these stress-adaptive pathways can be co-opted to support invasion, metastasis, and treatment resistance.

Our group previously identified an ASR gene signature associated with poor clinical outcomes across breast cancer subtypes, including involved in inflammatory signaling, oxidative stress adaptation, survival pathways, and protein translational regulation^15,18^. Importantly, subsets of this ASR metagene were associated with differences in survival outcomes between Black and White patients, suggesting that adaptation to stress responses may also contribute to biologic mechanisms underlying breast cancer disparities. This is significant as IBC is recognized as a breast cancer health disparity, with younger women and underrepresented populations experiencing disproportionately higher incidence rates, more aggressive disease, and worse survival outcomes^19–21^. Black patients with IBC are also more likely to present with advanced-stage disease and triple-negative tumors^22^, both of which are associated with inferior prognosis. Although disparities in access to care and structural inequities are important contributors to these outcomes, biologic mechanisms that may promote aggressive disease behavior in IBC remain incompletely defined. Because inflammatory and oxidative stress pathways are linked to both tumor aggressiveness and chronic systemic stress^23,24^, we postulate that the adaptive stress response signaling may represent a biologic mechanism connecting tumor emboli survival with clinical disparities in IBC.

Due to the rarity of IBC and limited availability of patient biospecimens, few studies have directly investigated the molecular programs that support tumor emboli survival. To address this gap, we used complementary preclinical tumor emboli models and patient transcriptomic datasets to define ASR programs specific to the tumor emboli phenotype in IBC. We identified an ASR-associated tumor emboli signature enriched for inflammatory, oxidative stress, and survival pathways and evaluated its association with clinicopathologic features, LVI, response to neoadjuvant therapy, and race-associated survival differences. Collectively, these findings establish adaptive stress signaling as a biologic framework linking tumor emboli survival, therapeutic resistance, and disparities in inflammatory breast cancer.

## Results

### Adaptive stress response genes are enriched in inflammatory breast cancer tumor emboli models

Tumor emboli within dermal lymphatic vessels are a defining pathologic feature of IBC (**Figure 1a**). To investigate whether ASR signaling contributes to IBC tumor emboli survival within the oxidative, inflammatory, and immune stress-rich lymphovascular microenvironment, we first evaluated expression of a previously defined set of ASR genes (ASR-226 metagene)^15^ (**Figure 1, middle panel**). We specifically examined whether ASR-226 genes overlapped with a previously reported 76-gene IBC-specific signature^25^ identified within this consortium dataset. Notably, 53 ASR-associated genes were present within the 76-gene IBC signature, suggesting that stress-adaptive signaling represents a prominent component of IBC biology (**Figure 1b)**. KEGG enrichment analysis of IBC-ASR-53 (**Supplementary Table 1a**) revealed WNT signaling, thyroid hormone synthesis and Bacterial invasion of epithelial cells as the top three molecular pathways, while white fat cell differentiation is the only significant biological process enriched by the overlapping gene set (**Supplementary Figure 1a-c** and **Supplementary Table 2a**). Gene interaction analysis using the STRING database further highlighted the interaction of genes associated with Immune and TGFβ signaling (**Figure 1c)**.

**Figure 1.**
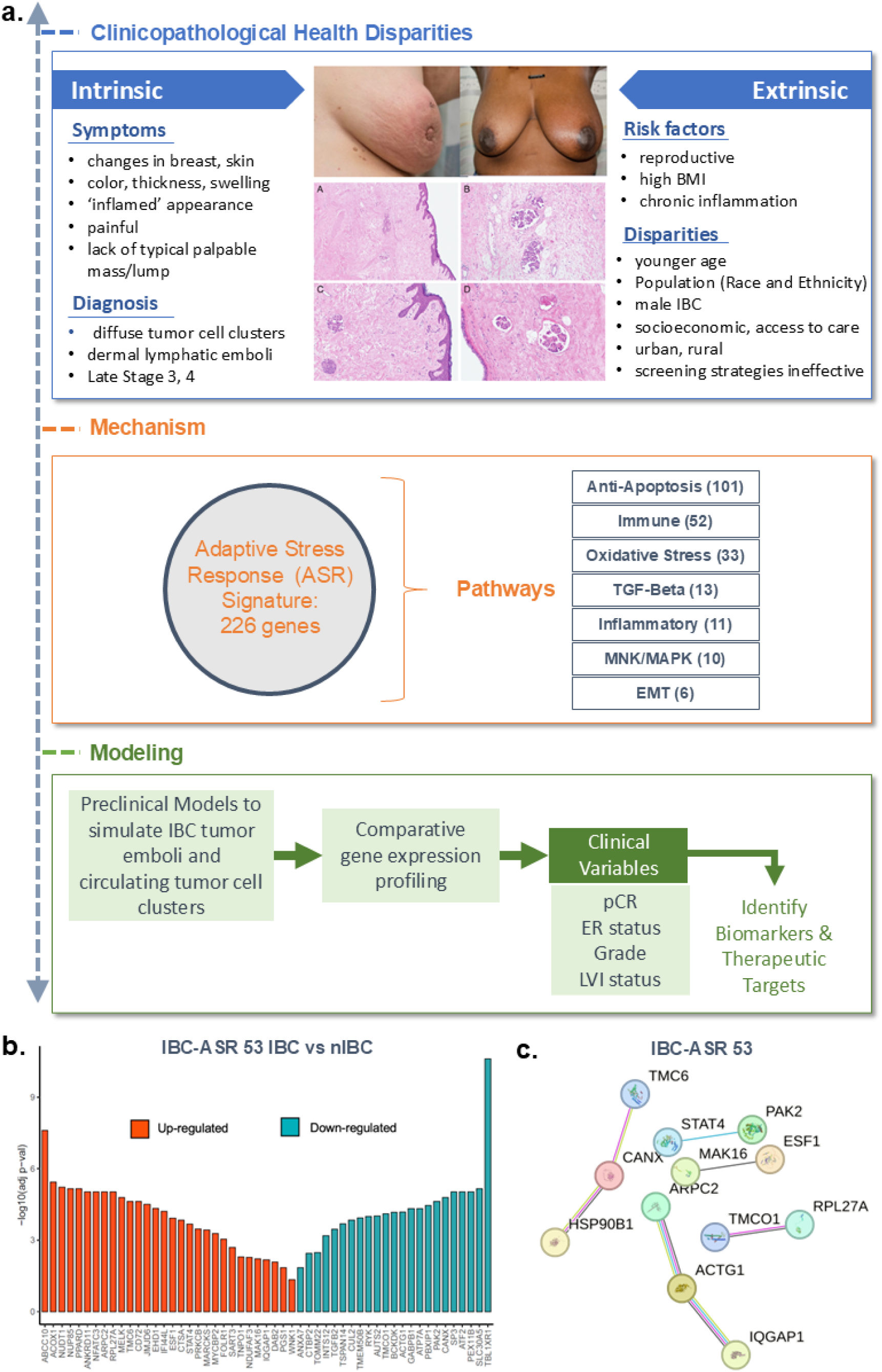
Inflammatory Breast Cancer Signature and Adaptive Stress Response Metagene. **a.** Schematic presentation of study design: Modelling Inflammatory Breast Cancer to study Adaptive Stress Response. **b.** Illustration of the overlap between Inflammatory Breast Cancer (IBC) gene signature and Adaptive Stress Response (ASR) gene set in IBC. The plot shows −log10 adjusted p value and the direction of change for each significant ASR gene in IBC vs. non-IBC samples. Red indicates upregulated genes and Blue indicates downregulated genes in IBC samples compared to non-IBC (adj.p-value <0.05) **c.** Visualization of the gene network using STRING interactome database for the overlapping genes between IBC gene signature and ASR gene set in IBC.

To further define functional ASR programs associated with the tumor emboli phenotype, we employed two complementary preclinical systems that recapitulate key features of IBC dissemination. The first model is a three-dimensional tumor–lymphatic architecture biomimetic platform (3D T-LAB)^14^ that allows isolation of circulating tumor cell clusters that simulate lymphatic invasion, termed TLAB-CC. The second model utilized tumor emboli-forming organoid cultures, termed TE. RNA sequencing of the TLAB-CC and TE cells derived using SUM149, a well-characterized pretreatment IBC patient-derived model widely regarded as one of the few true IBC-like cell lines^26^, identified distinct ASR-associated gene programs, including a 29-gene subset in the TLAB-CC compartment (ASR-29-TLAB-CC; **Figure 2a; Supplementary Table 3**) and a 28-gene subset in the tumor emboli organoid model (ASR-28-TE; **Figure 2a; Supplementary Table 4**).

**Figure 2.**
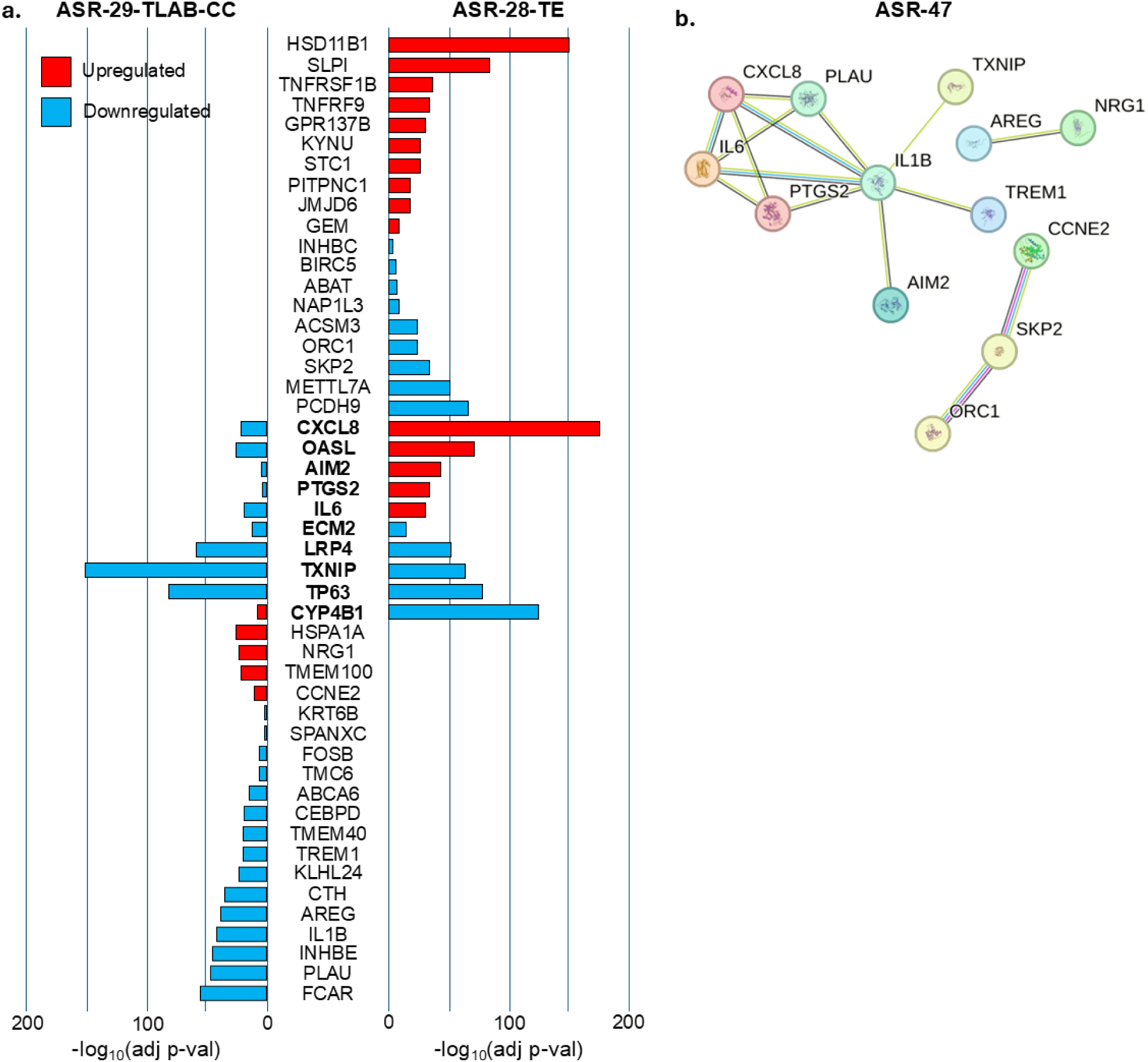
Identification of ASR gene sets in preclinical models of lymphatic circulating cells and IBC tumor emboli. **a**. Visualization of differentially expressed genes in the two models. The left panel of the plot shows −log10 adjusted p value and the direction of change for each significant ASR gene in TLAB-CC vs. 2D SUM149 samples. Red indicates upregulated genes; Blue indicates downregulated genes. 29/226 DEGs (termed ASR-29 T-LAB-CC gene set) that were statistically significant (adjusted p-value <0.05; log FC ≥ 2). The right panel of the plot shows −log10 adjusted p value and the direction of change for each significant ASR gene in SUM149 derived TE vs. parental monolayer samples. 28/226 DEGs (termed ASR-28-TE gene set) that were statistically significant (adjusted p-value <0.05; log FC ≥ 2). The overlapped genes between ASR-29 T-LAB-CC and ASR-28-TE are shown in bold. **b**. Gene interaction analysis using STRING database for the ASR-47 gene set, union of significant genes from the two models (ASR-29 T-LAB-CTCC, ASR-28-TE).

Ten differentially expressed genes (DEGs) were overlapped (*CXCL8/IL8*, *CYP4B1*, *PTGS2*, *IL6*, *TXNIP*, *AIM2*, *OASL*, *ECM2*, *LRP4*, and *TP63*) between the two complementary preclinical models of IBC tumor emboli biology (**Figure 2a**). Although these genes were common to both models, none demonstrated concordant upregulation. Among the shared genes, CYP4B1 uniquely exhibited opposite regulation between the two models, suggesting that metabolic adaptation may represent a key feature distinguishing stationary tumor cell clusters in the breast stroma from circulating lymphatic cell clusters. In contrast, *ECM2, LRP4, TXNIP, and TP63* were consistently downregulated across both platforms. Several inflammatory and immune-related genes, including *CXCL8/IL8*, *IL6*, *PTGS2*, *AIM2*, and *OASL*, were upregulated in TE-derived cells but downregulated in TLAB-CC cells, potentially reflecting microenvironment-dependent adaptations and immune-related changes accompanying lymphatic dissemination. Enrichment analysis of ASR-47 (**Supplementary Table 1b**) revealed IL-17 signaling, Cytokine-cytokine receptor interaction, NOD-like receptor signaling, and NF-kappa B signaling as the most significant signaling pathways and Gene Ontology analysis (**Supplementary Figure 2a-c** and **Supplementary Table 5a**) revealed steroid hormone, regulation of heat generation and heat generation as the top three significant biological processes. Despite these differences, their interaction analysis using the STRING database (**Figure 2b**) demonstrated convergence on common stress-adaptive mechanisms, with significant enrichment of XIAP-NFκB signaling, and oxidative stress response pathways.

### Tumor emboli-associated genes correlate with aggressive clinicopathologic features in IBC

To evaluate the clinical relevance of the ASR-associated TE and TLAB-CC genes, we examined gene expression patterns across multiple IBC patient cohorts, including estrogen receptor-positive (ER+) IBC, triple-negative IBC (TN-IBC; hormone receptor expression <10%), residual tumors obtained following neoadjuvant therapy and mastectomy (ypResidual), and publicly available World Consortium datasets comprising IBC (WC-IBC) and non-IBC (WC-nIBC) tumors (**Table 1**). Cohorts were analyzed independently to minimize potential batch effects.

**Table 1.**
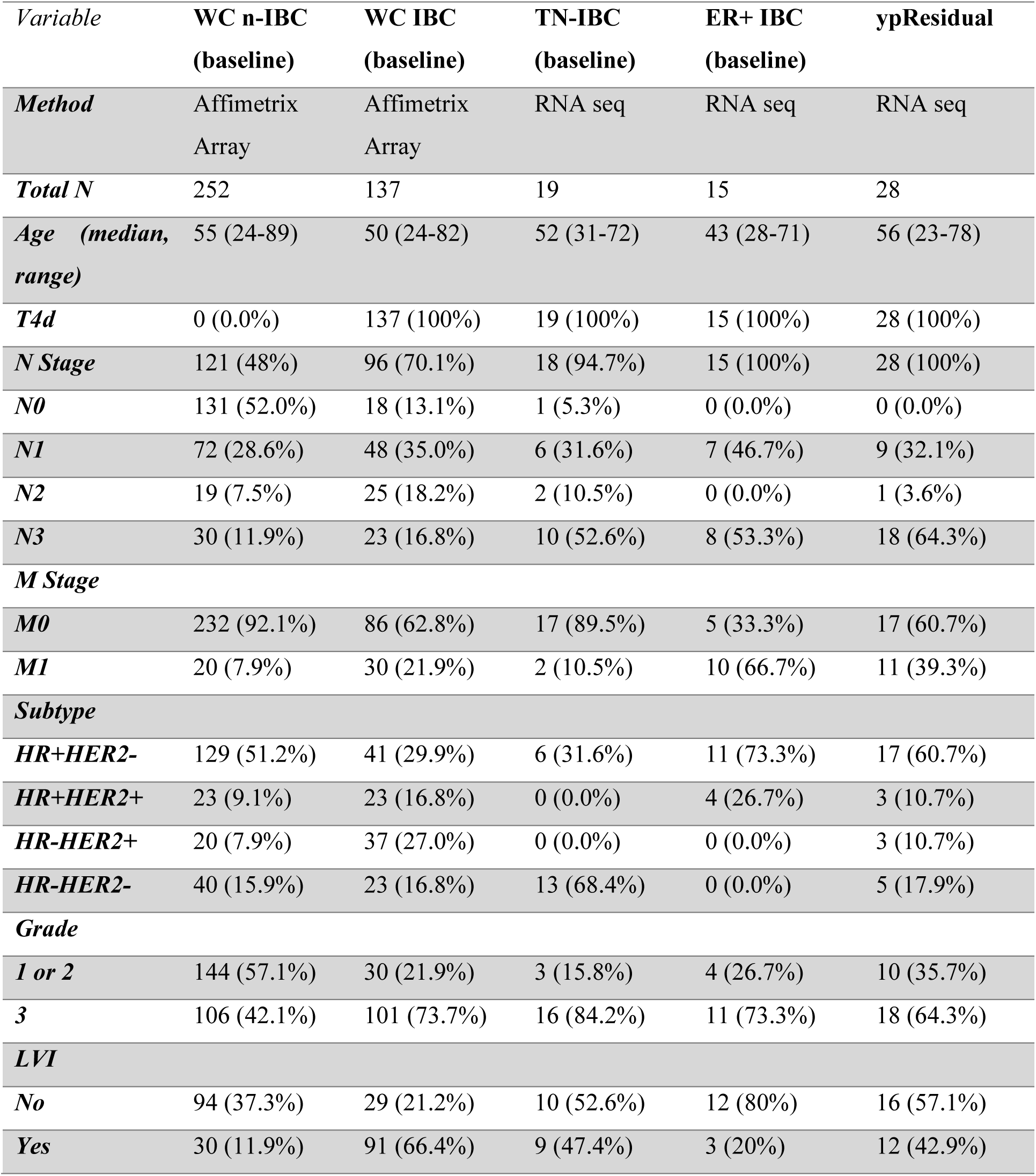
Clinicopathological parameters of the study cohort. WC-IBC: World consortium inflammatory breast cancer, WC nIBC: World consortium non-inflammatory breast cancer, ER+: Estrogen receptor positive, HR: Hormone receptor, HER: human epidermal growth factor receptor 2, LVI: Lymphovascular invasion.

| *Variable* | **WC n-IBC**<br>**(baseline)** | **WC IBC**<br>**(baseline)** | **TN-IBC**<br>**(baseline)** | **ER+ IBC**<br>**(baseline)** | **ypResidual** |
| --- | --- | --- | --- | --- | --- |
| **Method** | Affimetrix<br>Array | Affimetrix<br>Array | RNA seq | RNA seq | RNA seq |
| **Total N** | 252 | 137 | 19 | 15 | 28 |
| **Age (median, range)** | 55 (24-89) | 50 (24-82) | 52 (31-72) | 43 (28-71) | 56 (23-78) |
| **T4d** | 0 (0.0%) | 137 (100%) | 19 (100%) | 15 (100%) | 28 (100%) |
| **N Stage** | 121 (48%) | 96 (70.1%) | 18 (94.7%) | 15 (100%) | 28 (100%) |
| **N0** | 131 (52.0%) | 18 (13.1%) | 1 (5.3%) | 0 (0.0%) | 0 (0.0%) |
| **N1** | 72 (28.6%) | 48 (35.0%) | 6 (31.6%) | 7 (46.7%) | 9 (32.1%) |
| **N2** | 19 (7.5%) | 25 (18.2%) | 2 (10.5%) | 0 (0.0%) | 1 (3.6%) |
| **N3** | 30 (11.9%) | 23 (16.8%) | 10 (52.6%) | 8 (53.3%) | 18 (64.3%) |
| **M Stage** |  |  |  |  |  |
| **M0** | 232 (92.1%) | 86 (62.8%) | 17 (89.5%) | 5 (33.3%) | 17 (60.7%) |
| **M1** | 20 (7.9%) | 30 (21.9%) | 2 (10.5%) | 10 (66.7%) | 11 (39.3%) |
| **Subtype** |  |  |  |  |  |
| **HR+HER2-** | 129 (51.2%) | 41 (29.9%) | 6 (31.6%) | 11 (73.3%) | 17 (60.7%) |
| **HR+HER2+** | 23 (9.1%) | 23 (16.8%) | 0 (0.0%) | 4 (26.7%) | 3 (10.7%) |
| **HR-HER2+** | 20 (7.9%) | 37 (27.0%) | 0 (0.0%) | 0 (0.0%) | 3 (10.7%) |
| **HR-HER2-** | 40 (15.9%) | 23 (16.8%) | 13 (68.4%) | 0 (0.0%) | 5 (17.9%) |
| **Grade** |  |  |  |  |  |
| **1 or 2** | 144 (57.1%) | 30 (21.9%) | 3 (15.8%) | 4 (26.7%) | 10 (35.7%) |
| **3** | 106 (42.1%) | 101 (73.7%) | 16 (84.2%) | 11 (73.3%) | 18 (64.3%) |
| **LVI** |  |  |  |  |  |
| **No** | 94 (37.3%) | 29 (21.2%) | 10 (52.6%) | 12 (80%) | 16 (57.1%) |
| **Yes** | 30 (11.9%) | 91 (66.4%) | 9 (47.4%) | 3 (20%) | 12 (42.9%) |

Differential expression analyses identified distinct subsets of ASR-associated genes that were significantly associated with clinicopathologic variables. Because hormone receptor status is a major determinant of IBC biology, we first evaluated associations with ER expression. Analysis of the World Consortium datasets demonstrated that multiple ASR genes derived from the preclinical emboli models were significantly associated with ER status in both IBC and non-IBC tumors (**Figure 3a**). Several genes showed concordant expression patterns across WC-IBC, WC-nIBC, and ypResidual IBC tumors, suggesting that their associations were independent of endocrine therapy exposure. Specifically, *ABAT* and *CYP4B1* were upregulated in ER-positive tumors, whereas *HSD11B1*, *KYNU*, and *SLPI* were downregulated. Similar expression patterns were observed with progesterone receptor status, with additional downregulation of *CXCL8/IL8* and *SKP2* in PR-positive disease (**Supplementary Figure 3a**).

**Figure 3.**
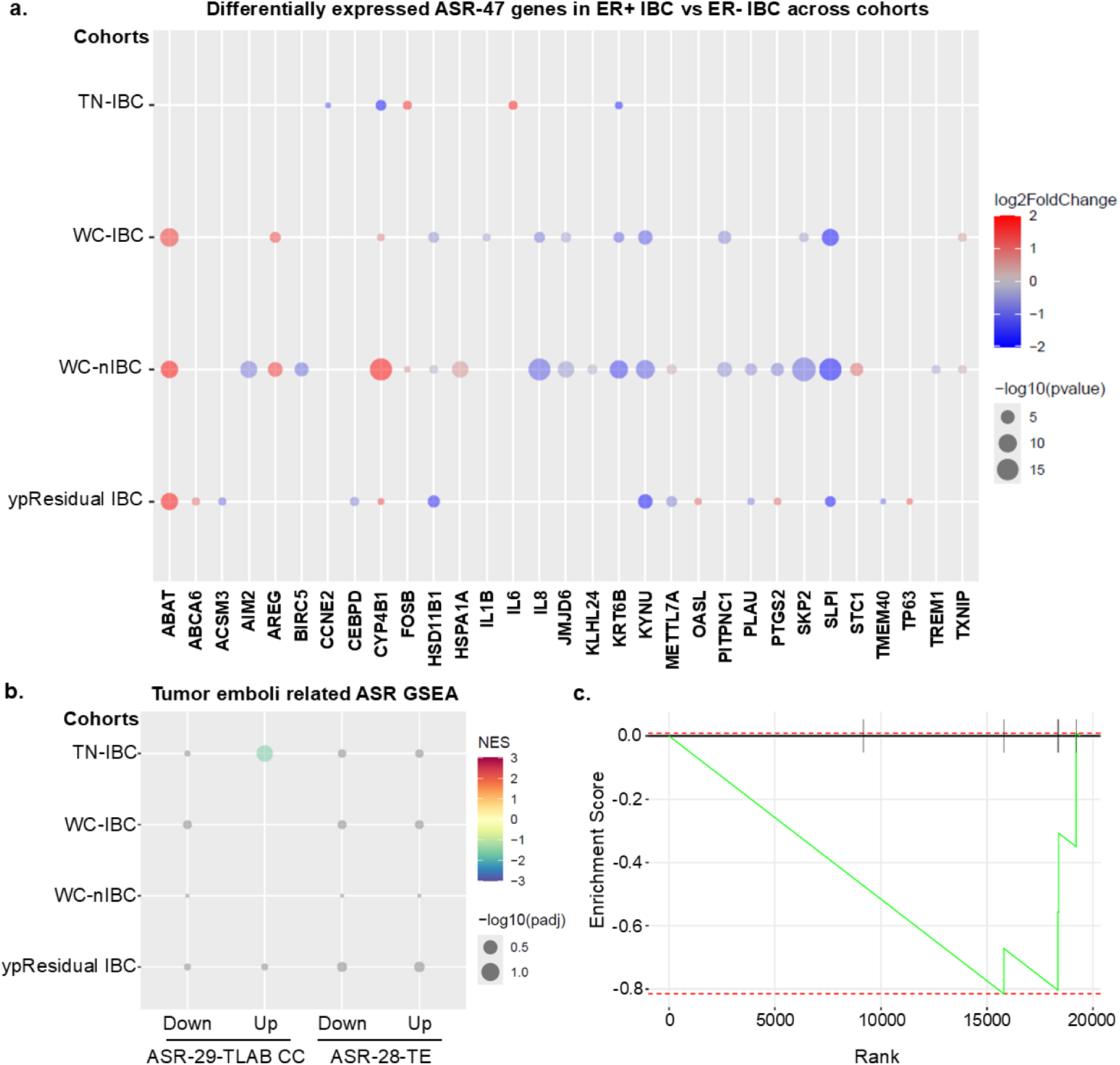
ASR-TE and ASR-TLAB-CC genes associated with Estrogen Receptor status in IBC and nIBC patient cohorts. **a.** Bubble plots show differentially expressed genes (DEGs, p-val < 0.05) for ER+ vs ER- in different cohorts where color indicates fold change and the size of the bubble represents the p-value. Genes are indicated on the X-axis and cohorts are indicated on the Y-axis. Sample size is as follows for TN-IBC cohort (Total n = 19; ER+ n = 15; ER- n = 4), WC-IBC cohort (Total n = 135; ER+ n = 67; ER- n = 68), WC-nIBC cohort (Total n = 250; ER+ n = 172; ER- n = 78) and ypResidual cohort (Total n = 28; ER+ n = 18; ER- n = 10). **b.** Bubble plot summarizing the enrichment analysis by ASR gene sets in different cohorts for ER+ vs ER- where color indicates Normalized Enrichment Score (NES) and the size of the bubble represents the p-value. Predefined gene sets are indicated on the X-axis and cohorts are indicated on the Y-axis **(left panel)**. **C.** Enrichment plot for significant ASR gene set ASR-29-TLAB-CC - Upregulated (adj p = 0.044937) for ER+ vs ER- patients in TN-IBC cohort **(right panel)**.

We next evaluated associations with tumor grade. Across ER+ and TN-IBC cohorts, *FOSB* and *SLPI* demonstrated concordant downregulation in high-grade tumors, whereas *OASL* expression was increased in TN-IBC and ypResidual tumors (**Figure 4a**). Gene set enrichment analysis (GSEA) of 4 signatures, ASR-28-TE – Upregulated, ASR-28-TE – Downregulated, ASR-29-TLAB-CC – Upregulated and ASR-29-TLAB-CC – Downregulated (details in methods) associated with ER status (**Supplementary Table 6a**), PR status (**Supplementary Table 6b**)., and tumor grade (**Supplementary Table 6c**) revealed distinct enrichment pattern linked to these clinical variables (**Figure 3b, c; Supplementary Figure 3b, c; Figure 4b-e**;).

**Figure 4.**
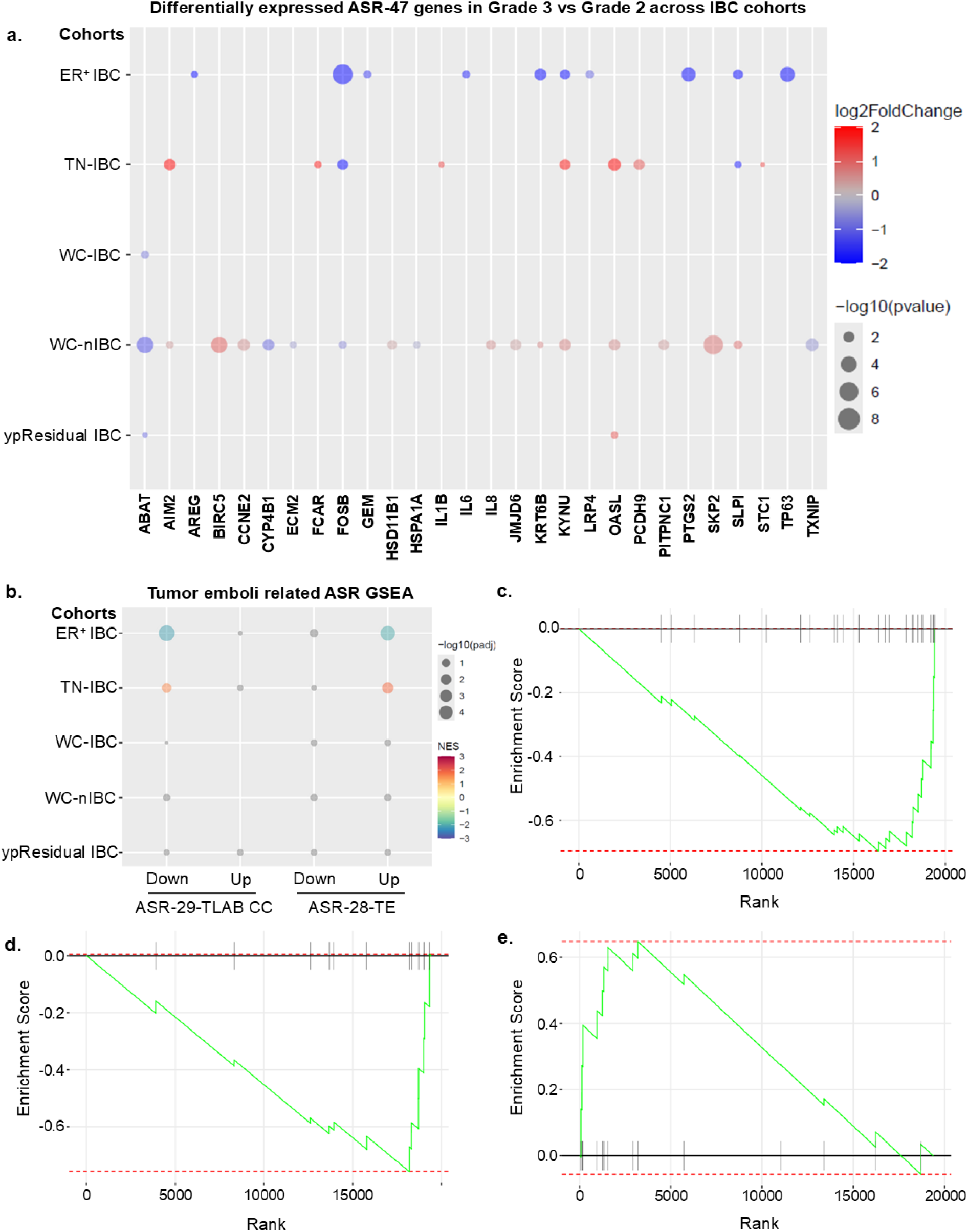
ASR-TE and ASR-TLAB-CC genes associated with Grade in IBC and nIBC patient cohorts. **a.** Bubble plots show differentially expressed genes (DEGs, p-val < 0.05) for Grade 3 vs. Grade2 in different cohorts where color indicates fold change and the size of the bubble represents the p-value. Genes are indicated on the X-axis and cohorts are indicated on the Y-axis. Sample size is as follows for ER+ cohort (Total n = 15; Grade 3 n = 11; Grade 2 n = 4), TN-IBC cohort (Total n = 19; Grade 3 n = 15; Grade 2 n = 4), WC-IBC cohort (Total n = 131; Grade 3 n = 101; Grade 2 n = 30), WC-nIBC cohort (Total n = 200 ; Grade 3 n = 106; Grade 2 n = 94) and ypResidual cohort (Total n = 28; Grade 3 n = 18; Grade 2 n = 10). **b.** Bubble plot summarizing the enrichment analysis by ASR gene sets in different cohorts for Grade 3 vs. Grade 2 where color indicates Normalized Enrichment Score (NES) and the size of the bubble represents the p-value. Predefined gene sets are indicated on the X-axis and cohorts are indicated on the Y-axis. Enrichment plots for significant ASR gene sets ASR-29-TLAB-CC - Downregulated **(c)** and ASR-28-TE – Upregulated **(d)** in ER+ cohort along with ASR-28-TE – Upregulated in TN-IBC **(e)** cohort (adj p < 0.05) for Grade 3 vs Grade 2 patients.

### Adaptive stress response genes are associated with lymphovascular invasion and dissemination phenotypes

Because LVI represents a critical step in tumor emboli dissemination and metastatic progression in IBC, we next examined whether the combined ASR-47 tumor emboli signature (ASR-28-TE and ASR-29-TLAB-CC) was associated with LVI status (**Figure 5**). In LVI-positive tumors within the ER+ cohort, *CTH*, *ECM2*, and *TNFSF9* were downregulated, whereas TMEM40 was upregulated. In contrast, LVI-positive TN-IBC tumors demonstrated upregulation of *CYP4B1*, *INHBC*, *CEBPD*, and *PCDH9*, accompanied by downregulation of *CXCL8/IL8* and *TNFSF9*. Additional associations involving *CEBPD*, *KYNU*, *SLPI*, and *SKP2* were observed in LVI-positive ypResidual tumors, linking ASR signaling to invasive and treatment-resistant disease states.

**Figure 5.**
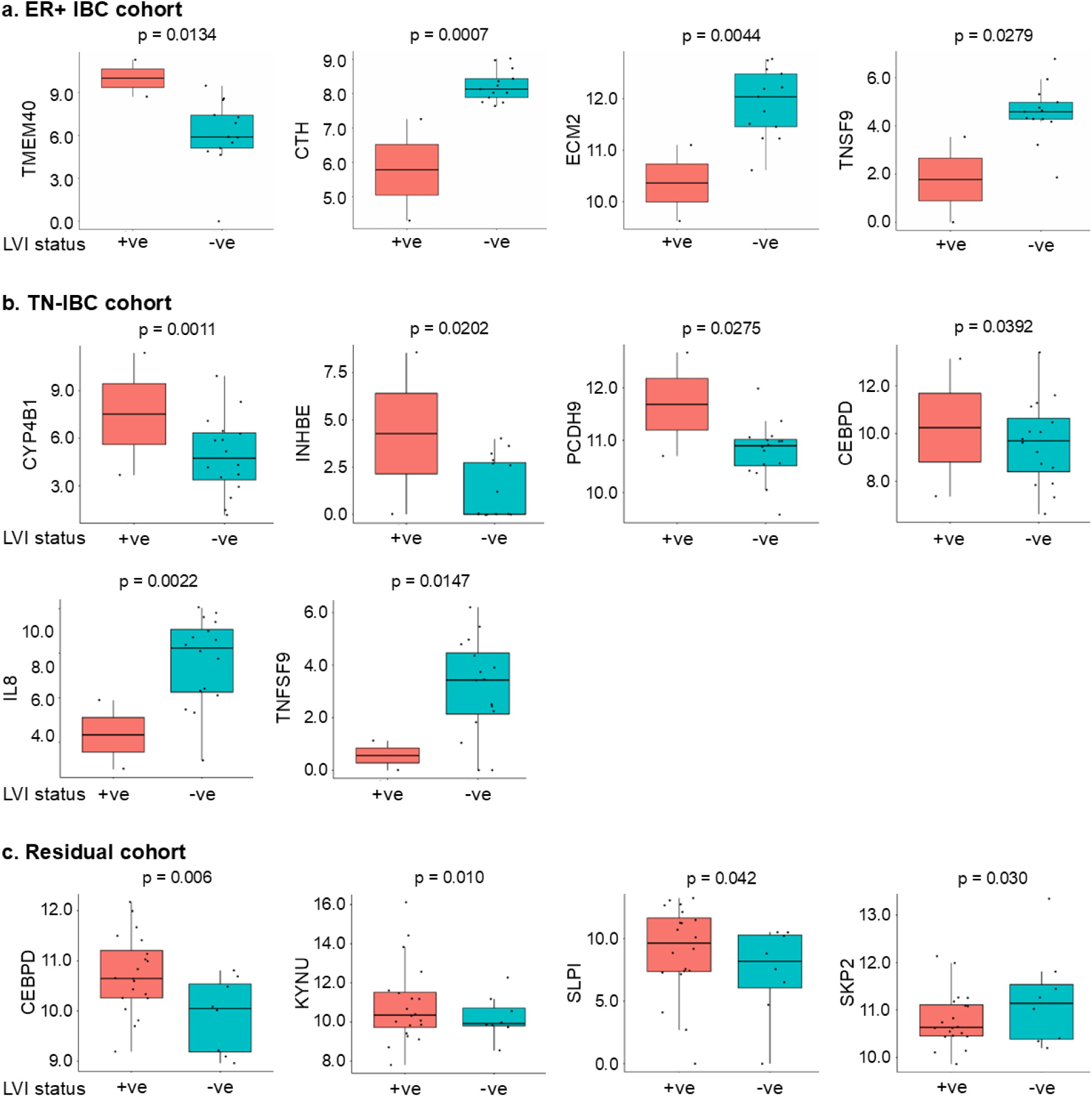
ASR genes associated with lymphovascular invasion in patient cohorts. Box plots summarizing gene expression in each patient for differentially expressed genes out of ASR-47 (p-val < 0.05) in three (ER+, TN-IBC and ypResidual) cohorts. **a.** In ER+ cohort (n = 15) for LVI +ve (n = 2) vs LVI -ve (n = 13), where *TMEM40* is upregulated and *CTH*, *ECM2*, *TNFSF9* are downregulated. **b.** In TN-IBC cohort (n = 18) for LVI +ve (n = 2) vs LVI-ve (n = 16), where *CYP4B1*, *INHBE*, *PCDH9*, *CEBPD* are upregulated and *IL8*, *TNFSF9* are downregulated. **c.** In ypResidual cohort (n = 28) for LVI +ve (n = 20) vs LVI -ve (n = 8), where *CEBPD*, *KYNU*, *SLPI* are upregulated and *SKP2* is downregulated. X-axis represents lymphovascular status where +ve stands for presence of LVI and –ve stands for absence of LVI and Y-axis represents gene expression. Dots represent corresponding gene expression values from individual patients.

To further evaluate the relationship between stress adaptation and lymphovascular dissemination, we interrogated the full ASR-226 metagene in the TN-IBC cohort. Four genes, *CYP4B1*, *STC5*, *IRF4*, and *SLC1A4*, were significantly associated with LVI presence (adjusted p<0.05; **Supplementary Table 7**). Notably, *CYP4B1* was also identified in both preclinical models and the ASR-226 metagene set, implicating metabolic stress adaptation as a potential mediator of tumor emboli survival and lymphatic dissemination.

Collectively, these findings support the postulate that adaptive stress signaling contributes not only to tumor emboli formation, but also to the invasive and lymphovascular dissemination phenotypes that characterize aggressive IBC biology.

### Adaptive stress response genes are linked to disparities in IBC outcomes

Given the disproportionate burden of IBC among Black women, we next investigated whether tumor emboli-associated ASR genes that we previously identified as being associated with clinical outcomes were differentially expressed between self-reported Black and White breast cancer patients^15^. To this end, we analyzed the combined ASR-47 tumor emboli signature (ASR-28-TE and ASR-29-TLAB-CC) in both the IBC cohorts described above and The Cancer Genome Atlas (TCGA) breast cancer cohort, a large publicly available dataset with diverse patient populations and long-term clinical outcomes. Analysis of the TCGA cohort identified that indeed self-reported black and white patients tend to cluster together within respective tumor subtypes (**Figure 6a**).

**Figure 6.**
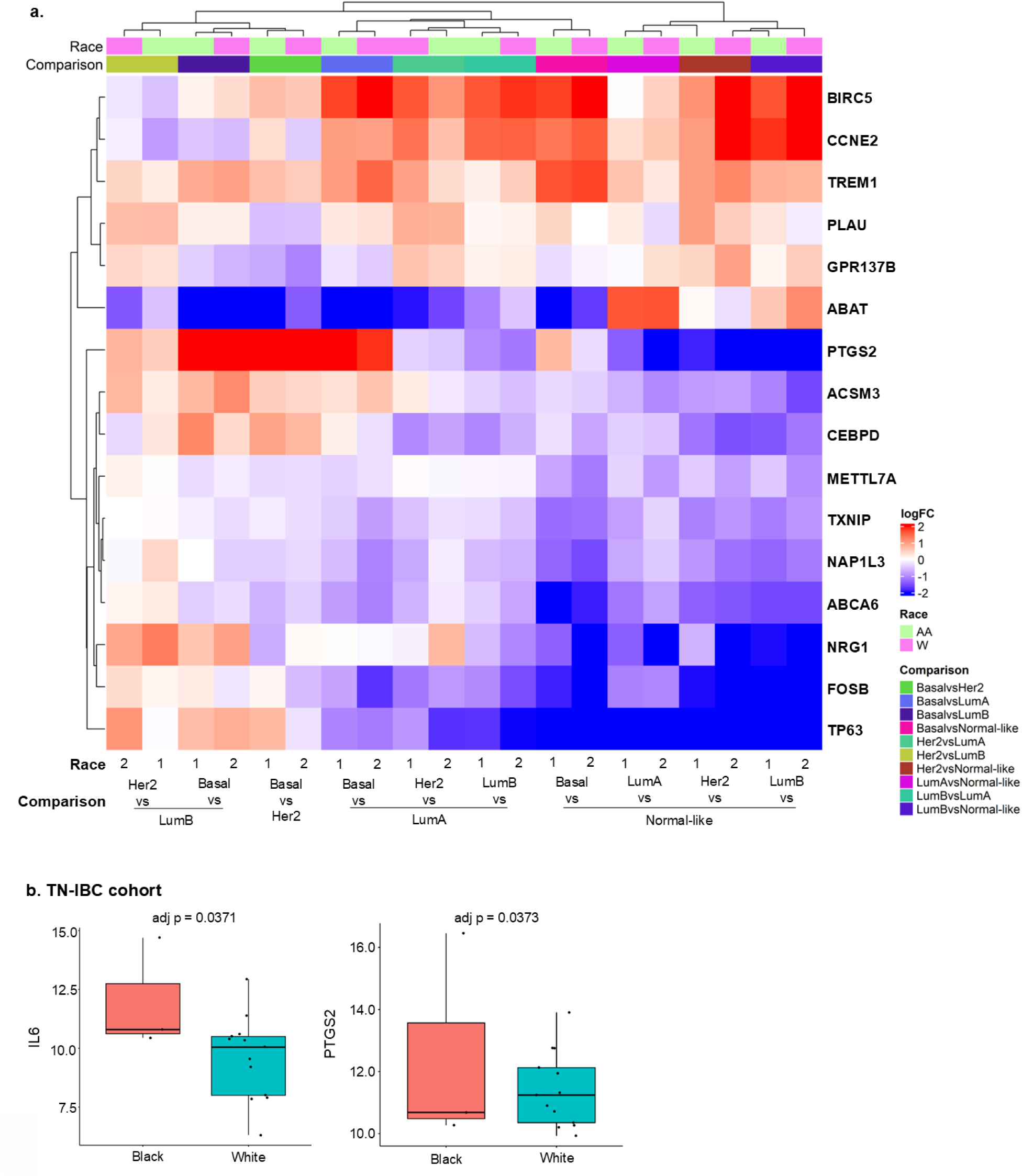
Gene expressions are associated with Black and White patients with breast cancer. **(a)** Heat map of 16 ASR genes associated with race related survival in different PAM 50 subtypes of breast cancer. Log fold change is shown in red, blue scale. X-axis, group 1 indicates Black breast cancer patients and group 2 indicates White breast cancer patients along with subtype comparison. **(b)** Box plots summarizing gene expression in each patient for differentially expressed genes out of ASR-47 (p-val < 0.05) in TN-IBC cohorts (n = 19) for Black patients (n = 3, Red) vs White patients (n = 13, Blue). X-axis represents patients and Y-axis represents gene expression. Each dot represents gene expression value for individual patients.

We next examined whether individual ASR-47 genes exhibited differential expression by race within the IBC cohorts. Notably, IL6 and PTGS2, both members of the ASR-47 tumor emboli program and identified in TCGA analyses, were significantly upregulated (adjusted p < 0.05) in Black patients with triple-negative IBC (TN-IBC) when compared with White patients (**Figure 6b**). Collectively, these findings implicate adaptive stress signaling as a potential biological contributor to disparities in IBC, linking inflammatory and survival pathways associated with tumor emboli biology to race-associated differences in disease outcomes.

### Adaptive stress response genes are associated with therapeutic response

Next, we evaluated associations between the ASR-47 genes and pathologic complete response (pCR) following neoadjuvant chemotherapy in cohorts with available treatment response data. Nine genes were significantly associated with pCR (**Figure 7**). *SKP2* and *AIM2* were enriched in tumors achieving pCR, whereas *CEBPD*, *CYP4B1*, *STC1, ABAT, BIRC5, and HSD11B1* were associated with residual disease and lack of pCR. *ECM2* exhibited cohort-specific behavior, showing increased expression in the TN-IBC cohort but decreased expression in the WC-IBC cohort. Functional annotation revealed that these genes mapped predominantly to oxidative stress adaptation and XIAP-NFκB survival pathways, suggesting that stress-adaptive programs contribute to therapeutic resistance and persistence of residual disease in IBC.

**Figure 7.**
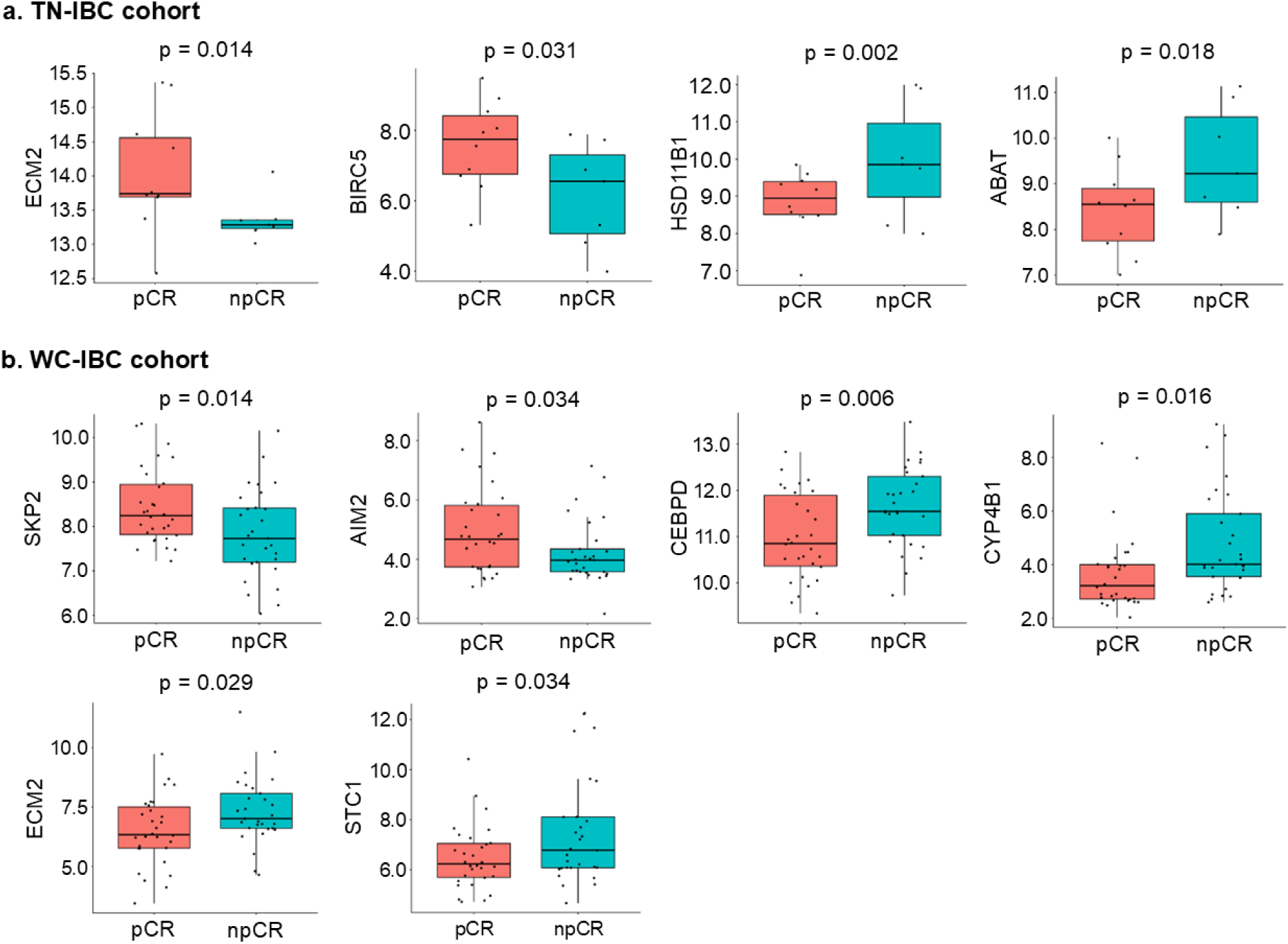
ASR and pathological complete response (pCR). Box plots summarizing gene expression in each patient for differentially expressed genes (DEGs) out of ASR-47 (p-val < 0.05). **a.** In TN-IBC cohort (n = 19) for pCR (n = 10) vs npCR (n = 7), where *ECM2* and *BIRC5* are upregulated whereas *HSD11B1* and *ABAT* are downregulated. **b.** In WC-IBC cohort (n = 59) for pCR (n = 30) vs npCR (n = 29), where *SKP2* and *AIM2* are upregulated whereas *CEBPD*, *CYP4B1*, *ECM2* and *STC1* are downregulated. X-axis represents pCR status and Y-axis represents gene expression. Each dot represents gene expression value for individual patient.

### Targeting adaptive stress response pathways suppresses IBC tumor emboli formation

To determine whether the ASR-associated tumor emboli signatures could be leveraged to identify therapeutic vulnerabilities, we functionally prioritized ASR genes associated with clinicopathologic variables, including lymphovascular invasion (LVI), according to their involvement in XIAP-NFκB survival signaling and oxidative stress response (OSR) pathways. Statistically significant ASR genes (p-val < 0.05) were grouped by functional pathway and average expression across cohorts (**Figure 8a; Supplementary Table 8**), revealing convergence of LVI-associated genes on these two stress-adaptive signaling networks.

**Figure 8.**
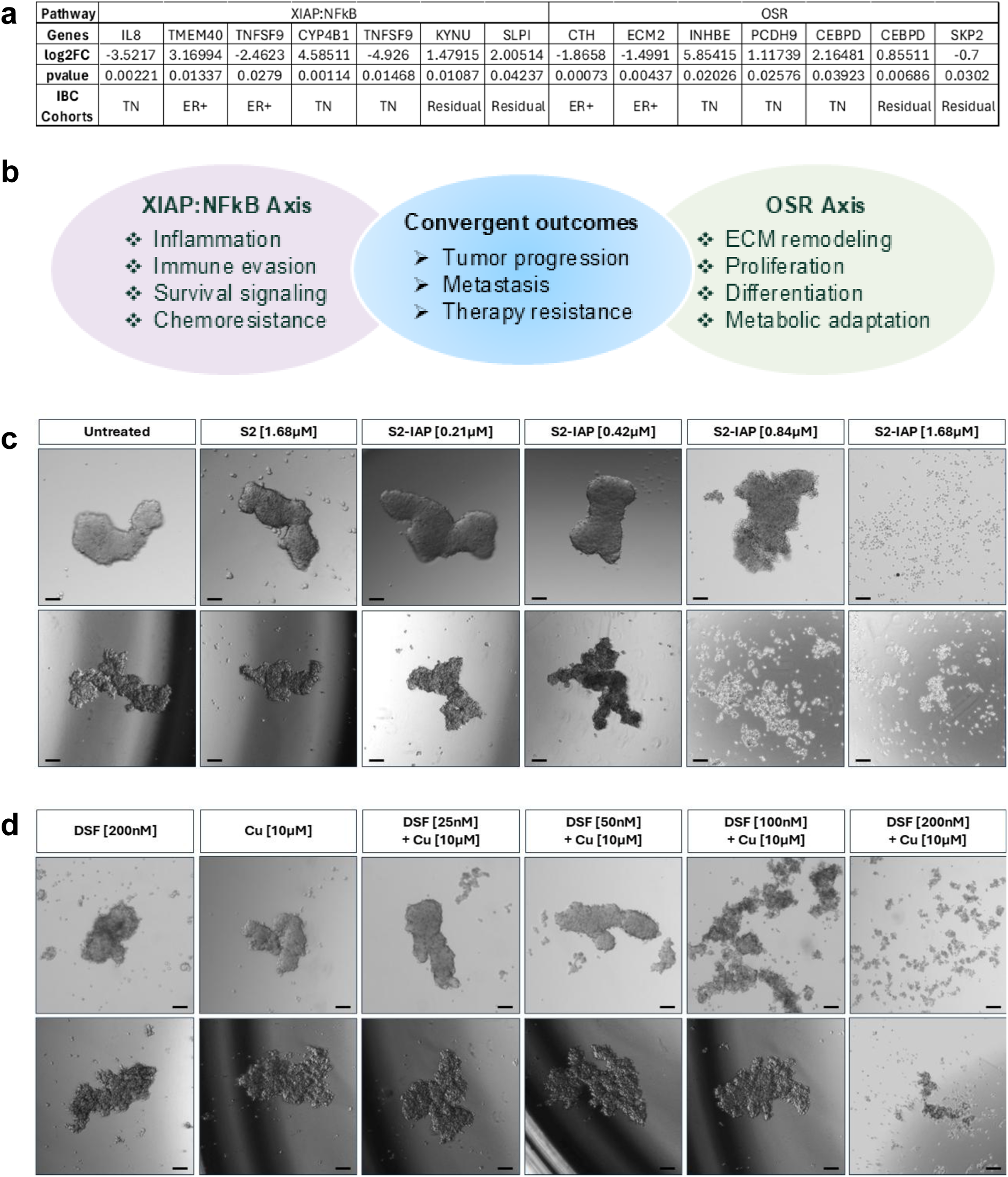
Targeting XIAP-NFkB and Oxidative Stress Response signaling pathways. **(a)** Top ASR-47 Genes that are significantly associated with LVI in patient cohorts are part of XIAP-NFkB or OSR signaling networks **(b)** Schematic representation of integrated OSR and XIAP signaling pathway highlighting their biological role. Effect of pharmacological inhibitors, S2-IAP **(c)** and DSF-Cu **(d)** on formation of tumor emboli derived from SUM149 basal-like (**top panel**) and MDA-IBC3 PDX **(bottom panel)**; Representative images at clinically relevant doses of at 24h post treatments are shown at the top of each column (n=3). Bar represents 100µm.

As proof of principle, we evaluated two pharmacologic agents with known activity against these pathways: the sigma-2 receptor-targeted SMAC mimetic S2-IAP^27,28^ and the oxidative stress response modulator disulfiram in combination with copper (DSF-Cu)^29,30^ (**Figure 8b**). Treatment with S2-IAP or DSF-Cu resulted in disruption of the tumor emboli formation as well as cell death in SUM149 cells and MDA-IBC3 patient-derived xenograft cells compared with vehicle-treated controls (**Figure 8 d, e; Supplementary figure 4**). Together, these results demonstrate that ASR-associated signaling pathways are not merely biomarkers of aggressive disease, but represent targetable dependencies required for survival of the tumor emboli phenotype.

## DISCUSSION

Herein, we report that adaptive stress response signaling represents a previously unrecognized feature of the tumor emboli phenotype in inflammatory breast cancer. Using complementary preclinical models and independent patient cohorts, we demonstrate that ASR-associated programs are enriched in tumor emboli and lymphatic circulating cell clusters and are associated with hormone receptor status, LVI, therapeutic response, and race-associated survival differences. Functional validation studies further identify XIAP-NFκB and oxidative stress response pathways as targetable dependencies capable of suppressing tumor emboli formation. Together, these findings establish a biologic framework linking stress adaptation to lymphovascular dissemination and suggest that ASR signaling represents a potential precision medicine axis in this highly aggressive breast cancer subtype.

Using complementary preclinical models of tumor emboli and lymphatic circulating cell clusters, we identified an ASR-associated signature enriched for XIAP-NFκB signaling, oxidative stress adaptation, inflammatory signaling, and immune regulatory pathways. These findings suggest that tumor emboli exist in a highly stress-adapted state that enables persistence during dissemination through inflammatory, oxidative, and immune stress-rich lymphovascular microenvironments. The distinct expression patterns observed between tumor emboli organoids and lymphatic circulating cell clusters further suggest that stress-adaptive programs are dynamic and evolve in response to microenvironmental pressures.

Validation in clinical cohorts identified concordant associations for *ABAT, CYP4B1, HSD11B1, KYNU, and SLPI* across World Consortium datasets and residual tumors following systemic therapy suggesting that these stress-adaptive programs reflect intrinsic tumor biology rather than treatment-induced effects. Importantly, our findings independently validate previous observations linking *ABAT* with ER-positive IBC ^31^ and extend these observations to aggressive non-IBC tumors, supporting the broader relevance of ASR signaling across breast cancer subtypes.

Another notable observation was the recurrent involvement of inflammatory mediators associated with the CXCL8-IL6-PTGS2 axis. *CXCL8* emerged as one of the most highly upregulated transcripts in the tumor emboli (TE) model and was shared between both preclinical platforms. However, *CXCL8*, together with *IL6* and *PTGS2*, was downregulated in lymphatic circulating cell clusters isolated from the T-LAB platform and in LVI-positive TN-IBC tumors, suggesting that inflammatory signaling programs may undergo dynamic remodeling during lymphatic dissemination. These findings suggest that tumor emboli may initially adopt a pro-inflammatory and pro-survival phenotype that subsequently transitions to a more immune-adapted state within the lymphovascular microenvironment. Recent studies, including in IBC, have implicated CXCL8 signaling in tumor plasticity, stemness, immune suppression, and therapeutic resistance across multiple malignancies, further supporting its role as a central mediator of stress adaptation^5,32,33^. Because CXCL8 is a well-established downstream target of XIAP-NFκB signaling^32^, our findings suggest that the CXCL8-IL6-PTGS2 inflammatory network may represent a critical component of the adaptive stress response underlying tumor emboli survival and dissemination in IBC.

One of the major findings of this study was the recurrent identification of genes involved in metabolic adaptation and oxidative stress regulation. Among these, CYP4B1 emerged as a particularly compelling candidate. *CYP4B1* was identified in both preclinical emboli models and was associated with ER status, LVI, and therapeutic response across multiple patient cohorts. Notably, *CYP4B1* was the only overlapping gene that demonstrated reciprocal expression between tumor emboli organoids and lymphatic circulating cell clusters, being increased within the 3DT-LAB compartment relative to tumor emboli/organoid cultures. Although the functional significance of this observation remains unknown, it raises the possibility that metabolic reprogramming may accompany lymphatic dissemination ^34^. Altered cytochrome P450 signaling has been implicated in oxidative stress tolerance and metabolic plasticity in aggressive malignancies^35^, suggesting that CYP4B1 may represent an important mediator of emboli survival.

Several additional genes repeatedly emerged across clinical and biologic contexts. *ECM2*, a gene abundantly expressed in adipose and mammary tissues ^36^, was consistently downregulated in both emboli-derived ASR subsets and associated with treatment response and LVI. *SLPI* correlated with hormone receptor status, tumor grade, and LVI and has previously been implicated in endothelial-coated tumor emboli formation^37^. Together with recurrent observations involving *HSD11B1, KYNU, CXCL8/IL8, SKP2, FOSB, OASL, TNFSF9, PTGS2, and CEBPD*, these findings suggest that extracellular matrix remodeling, inflammatory signaling, and metabolic adaptation represent previously underappreciated components of IBC biology that warrant further mechanistic investigation.

The association between ASR signaling and LVI, a defining feature of IBC and an essential step in metastatic dissemination is particularly noteworthy. We observed recurrent associations between ASR-emboli phenotype genes and LVI across ER-positive, triple-negative, and residual disease cohorts. Moreover, interrogation of the broader ASR-226 metagene identified *CYP4B1, STC5, IRF4,* and *SLC1A4* as additional genes associated with LVI in triple-negative IBC. These findings support a model in which stress-adaptive signaling programs contribute not only to tumor emboli formation but also to the invasive and dissemination phenotypes that characterize aggressive IBC.

A notable aspect of this study is the integration of stress-adaptive biology with cancer disparities. IBC disproportionately affects younger women and Black patients, who frequently experience more aggressive disease and poorer outcomes^38^. While structural inequities and differential access to care remain major drivers of these disparities, our findings suggest that tumor-intrinsic stress-adaptive pathways may also contribute to biologic mechanisms underlying aggressive disease behavior. *IL6* and *PTGS2* were significantly elevated in Black patients with TN-IBC, and multiple ASR-emboli phenotype genes were associated with race-related survival differences in breast cancer datasets. These findings are consistent with emerging evidence from our work and others that chronic inflammatory and oxidative stress pathways may intersect with social and environmental stress exposures to influence tumor biology and immune contexture^21,39–42,43–47^.

Importantly, the convergence of clinicopathologic correlates on XIAP-NFκB and oxidative stress response pathways enabled functional prioritization of therapeutic targets. Pharmacologic inhibition using a targeted SMAC mimetic (S2-IAP) and the FDA approved drug, Antabuse/disulfiram repurposed as an oxidative stress modulator in combination with copper^29,30^ resulted in marked suppression of tumor emboli formation in both SUM149 and MDA-IBC3 models. Both agents have previously been shown to disrupt stress-survival pathways implicated in tumor progression, metastasis, and therapeutic resistance in many cancer types^28,48^. These proof-of-concept studies suggest that ASR-associated pathways are not merely biomarkers of aggressive disease but may represent actionable dependencies required for maintenance of the tumor emboli phenotype.

We acknowledge several limitations in this study. The ASR-associated tumor emboli signature was derived from preclinical models and retrospective transcriptomic datasets, and prospective validation in independent IBC cohorts will be required. In addition, although genes such as *CYP4B1, SLPI, PTGS2, ECM2, and CEBPD* emerged recurrently across emboli formation, LVI, therapeutic response, and survival analyses, functional studies are needed to establish causal roles in dissemination and metastatic progression. Future studies incorporating spatial transcriptomics, single-cell analyses, and genetic or pharmacologic perturbation approaches may further elucidate how stress-adaptive signaling shapes the lymphovascular niche and therapeutic vulnerability in IBC.

In summary, we identify adaptive stress signaling as a previously underrecognized biologic feature of IBC and establish a framework linking stress adaptation to tumor emboli survival, lymphatic dissemination, therapeutic response, and disparities. These findings position ASR signaling as a potential precision medicine axis for biomarker development and therapeutic targeting in this rare and highly aggressive breast cancer subtype.

## METHODS

### Breast Cancer Samples and Expression Profiling

Gene expression data and clinicopathological data, including pathological response to neo-adjuvant chemotherapy, of patients with IBC and non-IBC were collected from publicly available affymetrix based gene expression datasets generated at Antwerp (E-MTAB-1006), Marseille (E-MTAB-1547), and Houston (E-GEOD-22597) as part of the Inflammatory Breast Cancer International Consortium (IBC-IC) as described previously^25^. Additional RNAseq based gene expression and patient clinicopathological data were obtained from the MD Anderson Cancer Center Morgan Welch Inflammatory Breast Cancer Clinic and Research Program. 19 TN-IBC^49^ patient baseline biopsies were from study (ClinicalTrials.gov Identifier: NCT02876107, registered on August 22, 2016). All had negative HER2 expression on immunohistochemistry (IHC) or fluorescence in situ hybridization (FISH) analysis; ER and PR IHC expression less than 10%. ER+ IBC – 35 baseline biopsy samples from confirmed IBC cases enrolled on the prospective IBC registry as above. 28 surgical samples (ypResidual) from confirmed IBC cases enrolled on the prospective IBC registry as above obtained after standard of care neoadjuvant systemic therapy. IBC specialists reviewed all cases in a prospective manner. Overall, patient inclusion criteria include histological confirmation of breast carcinoma, IBC confirmed according to international consensus criteria^33^. TCGA Research Network: https://www.cancer.gov/tcga is publicly available with prior patient’s consent and institutional review board agreements in place from original authors and the data related to breast cancer survival in Black and White patients previously published^15^ was queried for analysis of the ASR-emboli subsets.

### Preclinical Models and Expression Profiling

Gene expression data collected from RNA sequencing of SUM149-derived monolayer cells (n=3) tumor emboli cultures (TE) ^5,50^ (n=15) and circulating SUM149 cells (n=3) from 3D T-LAB biomimetic^14^ as described previously^18^. The identified ASR DEGs for TE vs SUM149 and TLAB-CC vs SUM149 are termed ASR-28-TE and ASR-29-TLAB-CC, respectively.

Briefly, RNA sequencing libraries were constructed utilizing the NEBNext Ultra II RNA Library Prep Kit for Illumina, adhering to the protocols outlined by the manufacturer (NEB, Ipswich, MA, USA). The quality of the sequencing libraries was assessed using the Agilent TapeStation (Agilent Technologies, Palo Alto, CA, USA) and quantified via the Qubit 2.0 Fluorometer (Invitrogen, Carlsbad, CA) and quantitative PCR (KAPA Biosystems, Wilmington, MA, USA). The libraries were then loaded onto a flowcell lane for clustering and the flowcell was placed into the Illumina HiSeq instrument (4000 or equivalent) for sequencing following the manufacturer’s guidelines. Sequencing was performed using a 2×150bp Paired End (PE) setup. Image processing and base calling were executed by the HiSeq Control Software (HCS). The raw sequence data in the form of .bcl files produced by the Illumina HiSeq was then converted into fastq files and de-multiplexed using the bcl2fastq 2.17 software from Illumina, allowing for one mismatch in the index sequence identification. Both library preparation and sequencing were conducted at Azenta US, Inc (South Plainfield, NJ, USA).

For T-LAB-CC, demultiplexed FASTQ sequencing reads were assessed using FastQC (v0.12.1) and MultiQC (v1.14)^51^, and adapter-contaminated and low-quality sequences were removed using Trimmomatic (v0.39)^52^. The trimmed reads were aligned to the human reference genome (GRCh38) using STAR (v2.7.10b)^53^. The GRCh38 reference genome were downloaded from GENCODE (v38.13)^54,55^. Normalization of sequence counts and differential expression analyses were performed using R (v4.2.2)^56^ and its extension package, DESeq2 (v1.38.2)^57^. ASR-29-TLAB-CC analyses were performed in R (version 4.2.2). To parse the microarray data into R, the following packages were utilized: hgu133plus2.db, jetset. We queried for the differentially expressed ASR metagene^15^ signature with a cut-off of Log2 fold change of 2 and an adjusted p-value of <0.05. ASR-28-TE RNA-Seq data were analyzed using Basepair software (https://www.basepairtech.com/) with a pipeline that included the following steps: (1) reads were aligned to the transcriptome derived from UCSC genome assembly hg19 using STAR 5 with default parameters; (2) read counts for each transcript were measured using feature counts 6; (3) differentially expressed genes were determined using DESeq2 7 and cut-off parameters of read count were 0.05 for adjusted P-value (corrected for multiple hypothesis testing). For T-LAB-CC vs. SUM149 monolayer, we used Benjamini–Hochberg (BH) approach.

### Gene Expression Analysis

Normalized expression values and raw read counts from preprocessed microarray and RNA-seq datasets are respectively utilized for downstream analyses of the five datasets (described above). Clinical comparisons were standardized as follows: Race (Black/African American vs. other), Grade (3 vs. 2), LVI (positive vs. negative), ER (positive vs. negative), PR (positive vs. negative), and pCR (pCR vs. non-pCR). Differential expression analysis was performed using the DESeq2 R package for RNA-seq data and two-sample t-tests for microarray data^57^. The resulting *p*-values from both platforms were adjusted for multiple testing using the Benjamini–Hochberg false discovery rate method^58^.. Genes with *p-val* < 0.05were considered differentially expressed. DEGs corresponding to adaptive stress response (ASR) signatures were visualized using dot plots, where dot color and size indicated direction and significance, respectively.

Functional evaluation of the transcriptomic data was conducted using a customized gene-set collection derived from previously established ASR gene signatures^15^. Pre-ranked GSEA was then performed using the test statistics obtained from the differential expression analyses across the clinical attributes of interest^59,60^. Functional annotation was performed using Gene Ontology (GO) enrichment analysis with the R package clusterProfiler^61^. Subsequently, network analysis of the gene sets of interest was conducted using the package rbioapi, leveraging the STRING functional interactome database^62,63^.

### Targeting Tumor emboli formation

SUM149 (IBC) was obtained from Asterand, Inc. and cultured as described previously^64^ and MDA-IBC3 (PDX) cells cultured as described previously^65^. Both cell lines were labeled with green fluorescent protein (GFP) as previously described^66^ and cultured according to established protocols^14,66,67^. For assessing inhibition of tumor emboli formation, we employed SUM149 and MDA-IBC3 derived emboli cultures as previously described^9,5^. Treatments were carried out at the time of seeding10mM stock solutions of DSF^29^, S2 and S2-IAP ^28^ were made in dimethyl sulfoxide (DMSO) and subsequently diluted in media to concentrations as indicated. 10mM stock of CuSO4 was made in the media. The highest final DMSO concentration of these solutions was below 0.02%. Phase contrast microscope images (4X) were taken on day 1 to assess tumor emboli formation using a EVOS M7000 (Thermo Ffisher Scientific) as described previously^5^.

## Data and Code availability

The datasets related to Gene expression used in this manuscript are available from the corresponding repositories without restrictions. Gene expression data (i.e., FASTQ-files, BAM-files, and associated metadata) of the 3D organoid of SUM149 have been deposited to ArrayExpress (E-MTAB-13929). Any other datasets used in this manuscript are available from the corresponding author on reasonable request.

## Author contributions

P.P., H.H., G.C.M., D.P.M.- methodology, analysis, validation, resources, writing, reviewing and editing. G.C.M., D.P.M., LD; S.V.L. - gene expression methodology, analysis, validation. G. R. D. - funding acquisition. W.G.H. preclinical. S.K. pathology. W.A.W., G.R.D.- conceptualization, methodology, analysis, validation, reviewing, editing, resources, writing-original draft preparation, editing, supervision. All authors have read and agreed to the published version of the manuscript.

## List of Abbreviations

3D T-LAB: 3D tumor-lymphatic architecture biomimetic
ASR: Adaptive Stress Response
ER: Estrogen Receptor
ER+ IBC: Estrogen Receptor positive Inflammatory breast cancer
GFP: Green Fluorescence Protein
IBC: Inflammatory Breast Cancer
IAP: Inhibitor of apoptosis family protein
LVI: Lymphovascular invasion
NFkB: Nuclear factor kappa-light-chain-enhancer of activated B cells
PDX: Patient derived xenograft
PR: Progesterone receptor
pCR: Pathological complete response
RFP: Red Fluorescence Protein
TE: Tumor Emboli
TLAB-CC: Circulating cells isolated from Tumor-Lymphatic Architecture Biomimetic
TN-IBC: Triple negative Inflammatory Breast Cancer cohort
WC-IBC: World Consortium-Inflammatory Breast Cancer cohort
WC-nIBC: World Consortium-non Inflammatory Breast Cancer cohort
XIAP: X-linked inhibitor of apoptosis

## Acknowledgements

The authors thank contributors from

The MDACC Inflammatory Breast Cancer (MDACC-IBC) Registry and DI Biopsy Team: Rachel Layman^1^, Bora Lim^1^, Azadeh Nasrazadani^1^, Sadia Saleem^1^, Vicente Valero^1^, Michael C. Stauder^2^, Wendy A. Woodward^2^, Anthony Lucci^3^, Susie X. Sun^3^, Gary J. Whitman^4^, Miral Patel^4^, Huong Le-Petross^4^, Yang Lu^4^, Angela Marx^1^, Angela Alexander^1^, Chasity Yajima^1^, Megumi Kai^1^, Lily Villareal^1^, Heather Lopez^1^.

1. Department of Breast Medical Oncology, Division of Cancer Medicine, The University of Texas MD Anderson Cancer Center, Houston, TX, USA
2. Department of Breast Radiation Oncology, Division of Radiation Oncology, The University of Texas MD Anderson Cancer Center, Houston, TX, USA
3. Department of Breast Surgical Oncology, Division of Radiation Oncology, The University of Texas MD Anderson Cancer Center, Houston, TX, USA
4. Department of Breast Imaging, Division of Diagnostic Imaging, The University of Texas MD Anderson Cancer Center, Houston, TX, USA

The World Consortium Dataset Leads from each represented institution: Naoto Ueno^1^, Francois Bertucci^2^, Steven Van Laere^3^,

1.University of Hawaii Cancer Ctr, Honolulu, HI

2 Predictive Oncology team, Centre de Recherche en Cancérologie de Marseille (CRCM), Inserm, CNRS, Aix-Marseille Université, Institut Paoli-Calmettes, Marseille, France

3. University of Antwerp, Belgium

Dr. Jing Wang, Department of Bioinformatics and Computational Biology, MD Anderson, Duke Inflammatory Breast Cancer Consortium at the Duke Cancer Institute; Dr. Kouros Owzar, Yanman Dai, Alexander Sibley at Duke Bioinformatics Shared resource; The Hollings Cancer Center CURE HUB (Cancer Research and Education Hub for Unified Biomedical, Clinical, and Population Science) team contributors: Hesham M. El-Shewy, Jack Whelchel, Zara Meaby for technical support for technical assistance and helpful discussions.

---

**Supplementary Figure 1.**
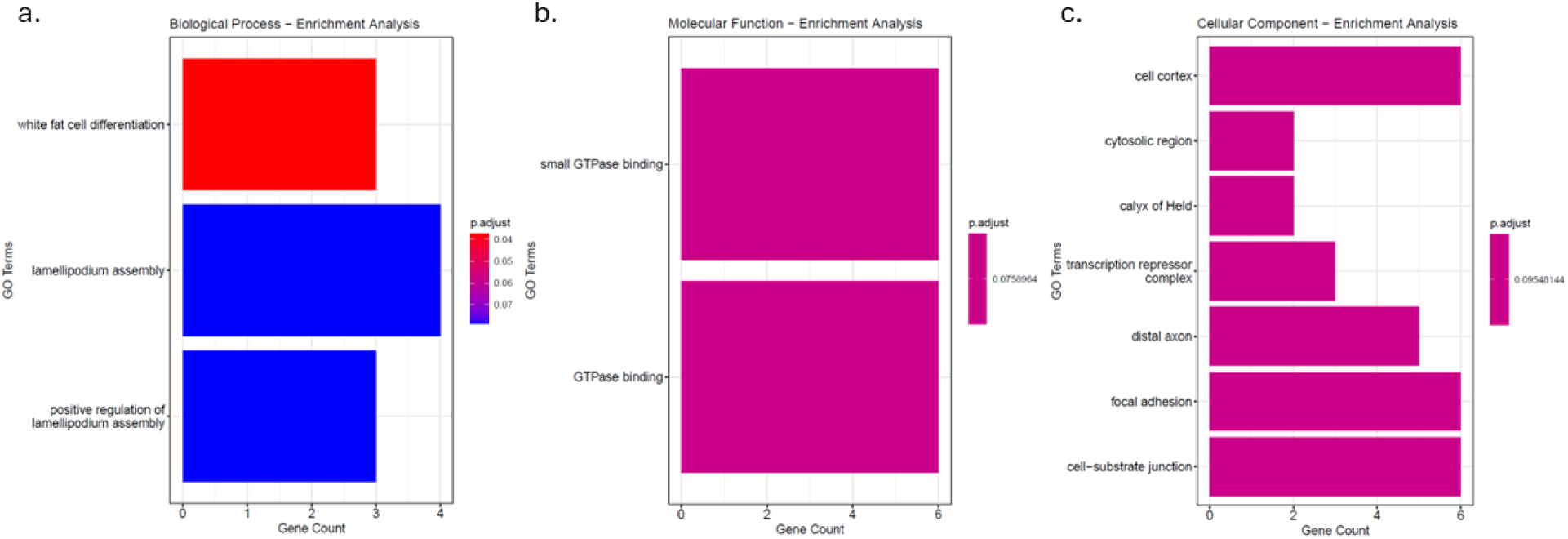
Gene ontology analysis IBC/ASR overlap genes. Statistically significant (p<0.05) Biological Process (**a**), Molecular Function (**b**) and Cellular Component (**c**) for overlapping 65 genes between Inflammatory Breast Cancer signature and Adaptive Stress Response metagene.

**Supplementary Figure 2.**
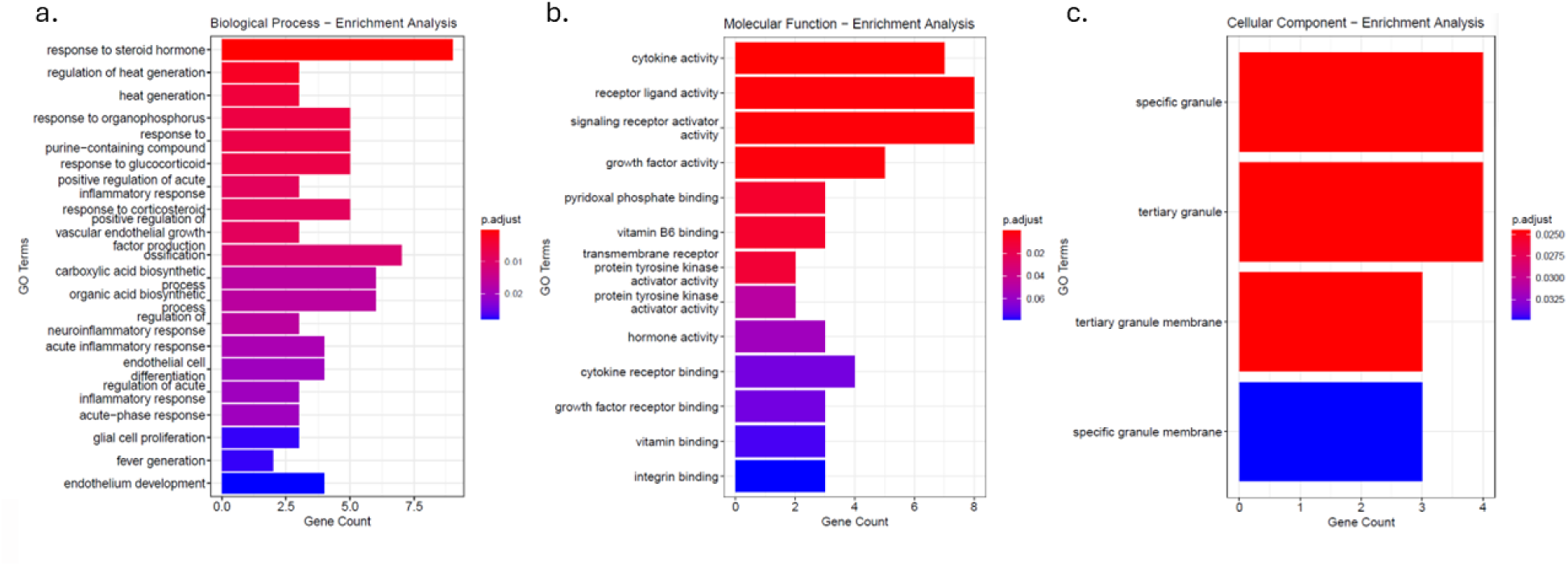
Gene ontology analysis for ASR-47. Statistically significant (p<0.05) Biological Process (**a**), Molecular Function (**b**) and Cellular Component (**c**) for ASR-47 (combined ASR-29-TLAB-CC and ASR-28-TE gene sets with 10 overlaps).

**Supplementary Figure 3.**
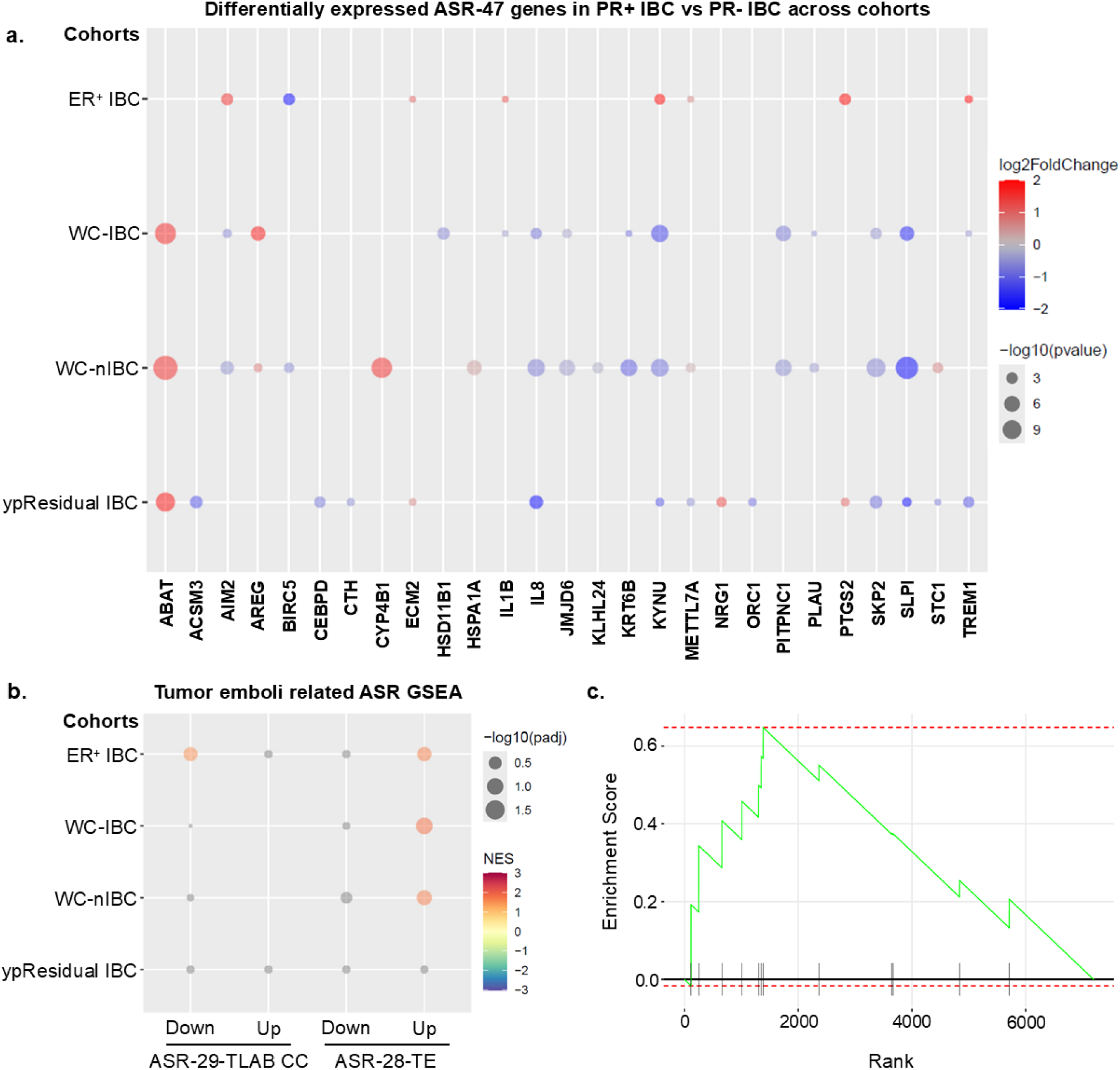
ASR-TE and ASR-TLAB-CC genes associated with Progesterone Receptor status in IBC and nIBC patient cohorts. **a.** Bubble plots show significantly differentially expressed genes (DEGs, p-val < 0.05) for PR+ vs PR-in different cohorts where color indicates fold change and the size of the bubble represents the p-value. Genes are indicated on the X-axis and cohorts are indicated on the Y-axis. Sample size is as follows for ER+ cohort (Total n = 15; PR+ n = 12; PR- n = 3), WC-IBC cohort (Total n = 135; PR+ n = 47; PR- n = 88), WC-nIBC cohort (Total n = 250; PR+ n = 138; PR- n = 112) and ypResidual cohort (Total n = 28; PR+ n = 15; PR- n = 13). **b.** Bubble plot summarizing the enrichment analysis by ASR gene sets in different cohorts for PR+ vs PR- where color indicates Normalized Enrichment Score (NES) and the size of the bubble represents the p-Value. Predefined gene sets are indicated on the X-axis and cohorts are indicated on the Y-axis **(left panel)**. **c.** Enrichment plot for significant ASR gene set ASR-28-TE - Upregulated in WC-IBC cohort (adj p < 0.05) **(right panel)**.

**Supplementary Figure 4.**
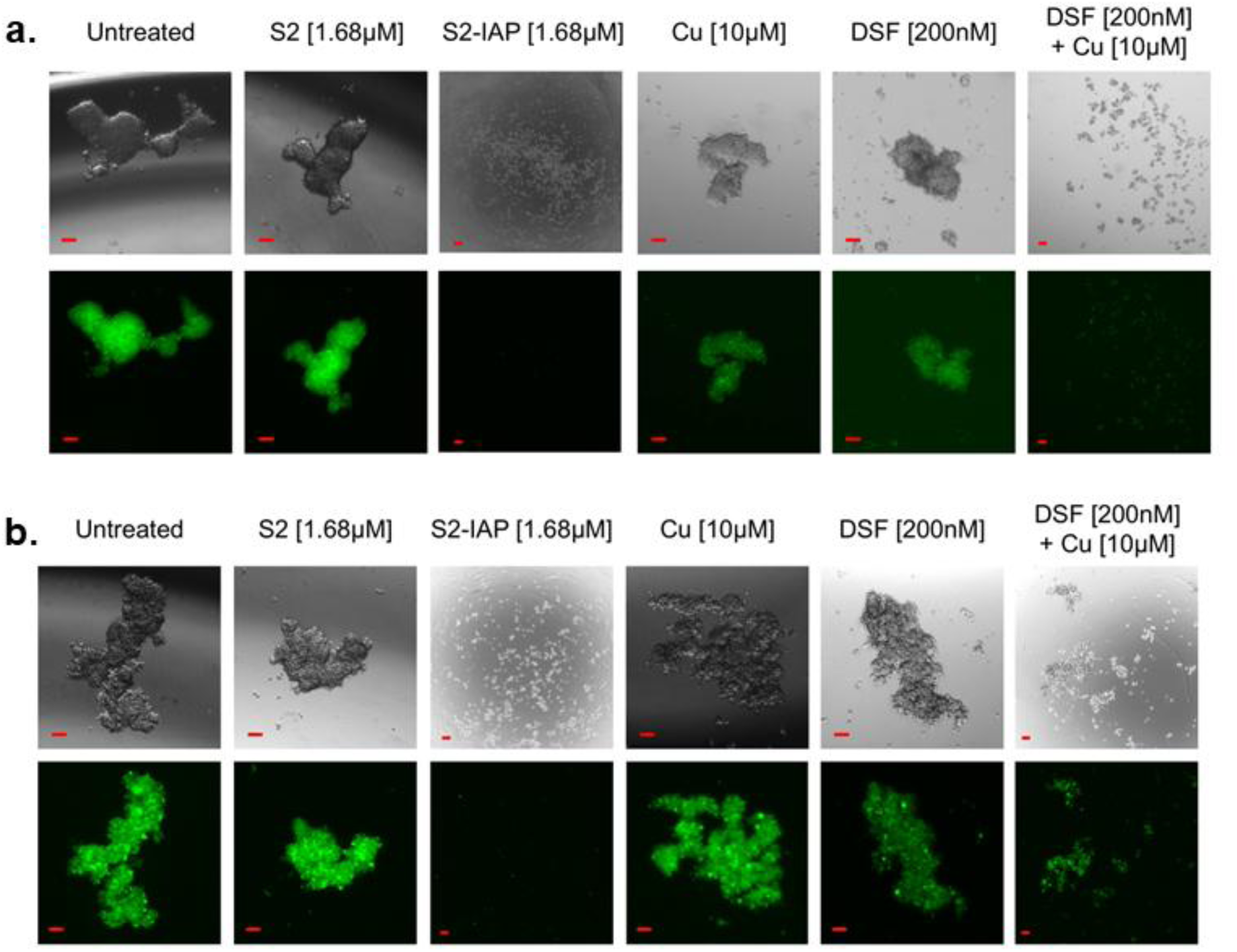
Inhibition of tumor emboli formation. SUM149 (**a**; GFP-tagged in green) and MDA-IBC3 (**b**; GFP-tagged in green) tumor emboli formation inhibition (n=3) via targeting OSR signaling pathway with DSF-Cu and XIAP signaling pathway with S2-IAP. Images are taken on Day 1 and treatments are shown on the top of each column. Bar represents 100µm.

**Supplementary Table 1. Enrichment analysis.** KEGG analysis for IBC-ASR-53 (**a**) and ASR-47 (**b**).

**Supplementary Table 2. Gene ontology analysis IBC/ASR overlap genes.** Statistically significant (p<0.05) Biological Process (**a**), Molecular Function (**b**) and Cellular Component (**c**) for overlapping 65 genes between Inflammatory Breast Cancer signature and Adaptive Stress Response metagene.

**Supplementary Table 3. ASR-29-TLAB-CC gene set.** Statistically significant (adj p<0.05) ASR genes, functional pathways, and expression TLAB-CC vs. 2D cell culture.

**Supplementary Table 4. ASR-28-TE gene set.** Statistically significant (adj p<0.05) ASR genes, functional pathways, and expression TE vs. 2D cell culture.

**Supplementary Table 5. Gene ontology analysis for ASR-47.** Statistically significant (p<0.05) Biological Process (**a**), Molecular Function (**b**) and Cellular Component (**c**) for ASR-47 (combined ASR-29-TLAB-CC and ASR-28-TE gene sets with 10 overlaps).

**Supplementary Table 6. Gene Set Enrichment Analysis (analysis) for four gene sets.** ASR-29-TLAB-CC and ASR-28-TE gene sets were subdivide based on expression directionality i.e., down- or up- regulation and these four gene sets were quired for enrichment in five cohorts (ER+ IBC, TN-IBC, WC-IBC, WC-nIBC and ypResidual Cohort) for ER+ vs ER- (**a**), PR+ vs PR- (**b**), Grade3 vs Grade 2 (**c**). Table shows fold change, significance and leading-edge genes.

**Supplementary Table 7.** Statistically significant genes associated with LVI presence (adjusted p<0.05) in TN-IBC cohort.

**Supplementary Table 8.** Statistically significant genes and their associated sub-ASR pathway correlated with LVI presence (p-val<0.05) in different cohorts.

